# Orthogonal chemical genomics approaches reveal genomic targets for increasing anaerobic chemical tolerance in *Zymomonas mobilis*

**DOI:** 10.1101/2025.07.09.663894

**Authors:** Jacob B. Eckmann, Amy L. Enright Steinberger, Morgan Davies, Elizabeth Whelan, Kevin S. Myers, Margaret L. Robinson, Amy B. Banta, Piyush B. Lal, Joshua J. Coon, Trey K. Sato, Patricia J. Kiley, Jason M. Peters

## Abstract

Genetically-engineered microbes have the potential to increase efficiency in the bioeconomy by overcoming growth-limiting production stress. Screens of gene perturbation libraries against production stressors can identify high-value engineering targets, but follow-up experiments needed to guard against false positives are slow and resource-intensive. In principle, the use of orthogonal gene perturbation approaches could increase recovery of true positives over false positives because the strengths of one technique compensate for the weaknesses of the other, but, in practice, two parallel screens are rarely performed at the genome-scale. Here, we screen genome-scale CRISPRi (CRISPR interference) knockdown and TnSeq (transposon insertion sequencing) libraries of the bioenergy-relevant Alphaproteobacterium, *Zymomonas mobilis*, against growth inhibitors commonly found in deconstructed plant material. Integrating data from the two gene perturbation techniques, we established an approach for defining engineering targets with high specificity. This allowed us to identify all known genes in the cytochrome *bc*_1_ and cytochrome *c* synthesis pathway as potential targets for engineering resistance to phenolic acids under anaerobic conditions, a subset of which we validated using precise gene deletions. Strikingly, this finding is specific to the cytochrome *bc*_1_ and cytochrome *c* pathway and does not extend to other branches of the electron transport chain. We further show that exposure of *Z. mobilis* to ferulic acid causes substantial remodeling of the cell envelope proteome, as well as the downregulation of TonB-dependent transporters. Our work provides a generalizable strategy for identifying high-value engineering targets from gene perturbation screens that is broadly applicable.

**IMPORTANCE:** Engineering microorganisms to tolerate harsh production conditions stands to increase bioproduct yields of engineered microbes. In this study, we systematically identified *Z. mobilis* genes that confer resistance or susceptibility to chemical stressors found in deconstructed plant material. We used complementary genetic techniques to cross-validate these genes at scale, providing a widely applicable method for precisely identifying genetic alterations that increase chemical resilience. We discovered genetic modifications that improve anaerobic growth of *Z. mobilis* in the presence of inhibitory chemicals found in renewable plant-based feedstocks. These results have implications in engineering robust production strains to support efficient and resilient bioproduction. Our methodologies can be broadly applied to understand microbial responses to chemicals across systems, paving the way for developments in biomanufacturing, therapeutics, and agriculture.

## INTRODUCTION

Industrial-scale fermentation by genetically engineered microorganisms can produce a wide array of products, including fuels and chemicals traditionally derived from petroleum. Over the last several decades, renewable plant biomass has emerged as a promising carbon source for conversion to value-added bioproducts (1,2). However, plant-based feedstocks also contain a myriad of known and unknown chemical stressors which slow microbial growth and limit fermentation yields (3–5). These chemicals are derived from breakdown of lignocellulose from plant cell walls (e.g., aromatic and phenolic acids, furans), solvents used to unlock sugars from biomass (e.g., ammonia or γ-valerolactone) or fermentation products themselves (e.g., ethanol and isobutanol) and can affect microorganisms through a broad array of mechanisms including changing membrane permeability and intracellular pH, generating reactive oxygen species (ROS), interacting with and mutagenizing DNA, and directly inhibiting cellular enzymes (5–11). Engineering microorganisms to better tolerate these chemical stressors stands to increase fermentation yields.

One way to identify genetic engineering targets to improve resilience is through chemical genomics, where mutant libraries are screened against a suite of inhibitory chemicals to reveal changes in mutant fitness resulting from chemical exposure (12–15). As a high-throughput assay, chemical genomics is subject to false positives that can expend time and resources to follow-up if not avoided. Parallel use of multiple orthogonal mutant libraries could theoretically reduce false positives through cross-validation of phenotypes, though this approach is underexplored.

TnSeq and CRISPRi libraries are ideal tools for this comparative approach, as they have complementary strengths and weaknesses (16). For example, screening TnSeq libraries can identify essential genes but cannot further characterize their chemical phenotypes because transposon insertion in an essential gene is lethal by definition. CRISPRi libraries can be designed to overcome this limitation by including mismatched single guide (sg)RNAs for partial knockdown of essential genes (Mismatch-CRISPRi) or by partially inducing CRISPRi components, maintaining cell viability and allowing for phenotype identification (17). Additionally, the two technologies cause different *cis* effects on downstream gene expression, potentially leading to distinct types of false positives. CRISPRi reliably reduces expression of the gene target and all downstream genes in a transcription unit, a phenomenon called polarity (18,19). Conversely, transposons often contain a strong promoter to drive expression of a selective marker, and transcription across the insertion junction can result in readthrough expression of downstream genes. This has been used as a strategy for deliberate screening of gene overexpression phenotypes (20). Therefore, integration of both approaches could reduce false positives by identifying targets with consistent phenotypes despite differing *cis* effects.

The Alphaproteobacterium *Zymomonas mobilis*, a facultative anaerobe well known for its native ability to rapidly convert sugars into ethanol, is a promising organism for industrial fermentation of renewable plant biomass to bioproducts (21–24). Enhancing its industrial appeal, *Z. mobilis* features high native tolerance to ethanol and some, but not all, lignocellulose-derived chemicals (25,26). It also features a small genome with an improving toolkit for genomic manipulation (26–29). However, much of the *Z. mobilis* genome remains unannotated, impeding rational engineering efforts to further enhance stress tolerance. To address this challenge, other groups have performed chemical genomics on *Z. mobilis* Tn libraries grown aerobically in bioenergy-relevant inhibitors using a microarray-based quantification approach that was a predecessor to TnSeq (15, 30). However, large batch industrial fermentations with *Z. mobilis* are likely to be grown anaerobically given that *i*) the optimal growth environment of *Z. mobilis* is anaerobic, and *ii*) the high cost of aerating industrial-scale fermenters. Thus, deciphering how *Z. mobilis* responds to inhibitory production stressors under anaerobic conditions remains an important question.

In this study, we use TnSeq and CRISPRi as orthogonal approaches to identify genes that alter susceptibility to fermentation-relevant chemicals in *Z. mobilis* under anaerobic conditions. We quantitatively compare the two techniques to increase reliability of discoveries by enriching for true positives. Our screens reveal a surprising role for an electron transport pathway in anaerobic chemical stress caused by phenolic acids (e.g., ferulic acid) encountered during microbial fermentation of plant biomass, providing genetic engineering targets for increasing microbial growth and bioproduct yields. Finally, we characterize the effect of ferulic acid on the *Z. mobilis* proteome, revealing potential mechanisms of phenolic acid-induced stress.

## RESULTS

### Parallel screening detects genes that alter fitness against production stressors

To identify *Z. mobilis* genes that modulate fitness in the presence of industrially-relevant chemicals, we screened TnSeq and CRISPRi libraries anaerobically against a suite of inhibitory production stressors at sub-lethal concentrations (Fig. S1, Table S1). For each gene knockout (TnSeq) or knockdown (CRISPRi), we calculated a chemical-gene (CG) score representing the log_2_ fold change in relative mutant abundance between chemical-treated and untreated cultures. Positive CG scores indicate mutants which have increased fitness with the chemical, while negative CG scores indicate mutants that have decreased fitness.

Theorizing that parallel use of orthogonal techniques could increase reliability of results, we developed a method for cross-validating findings between the two libraries (Fig. 1A). Various technical factors, such as sequencing depth, library size, and population doublings, may impact the spread of CG scores for each library in each condition. To account for these factors, we first quantile normalized the CG scores for non-essential genes (i.e., genes which could be assessed by both the TnSeq and CRISPRi libraries) for both libraries within each condition, producing consistent distributions of CG scores that facilitates direct comparison of the data sets (Fig S2).

**Figure 1.**
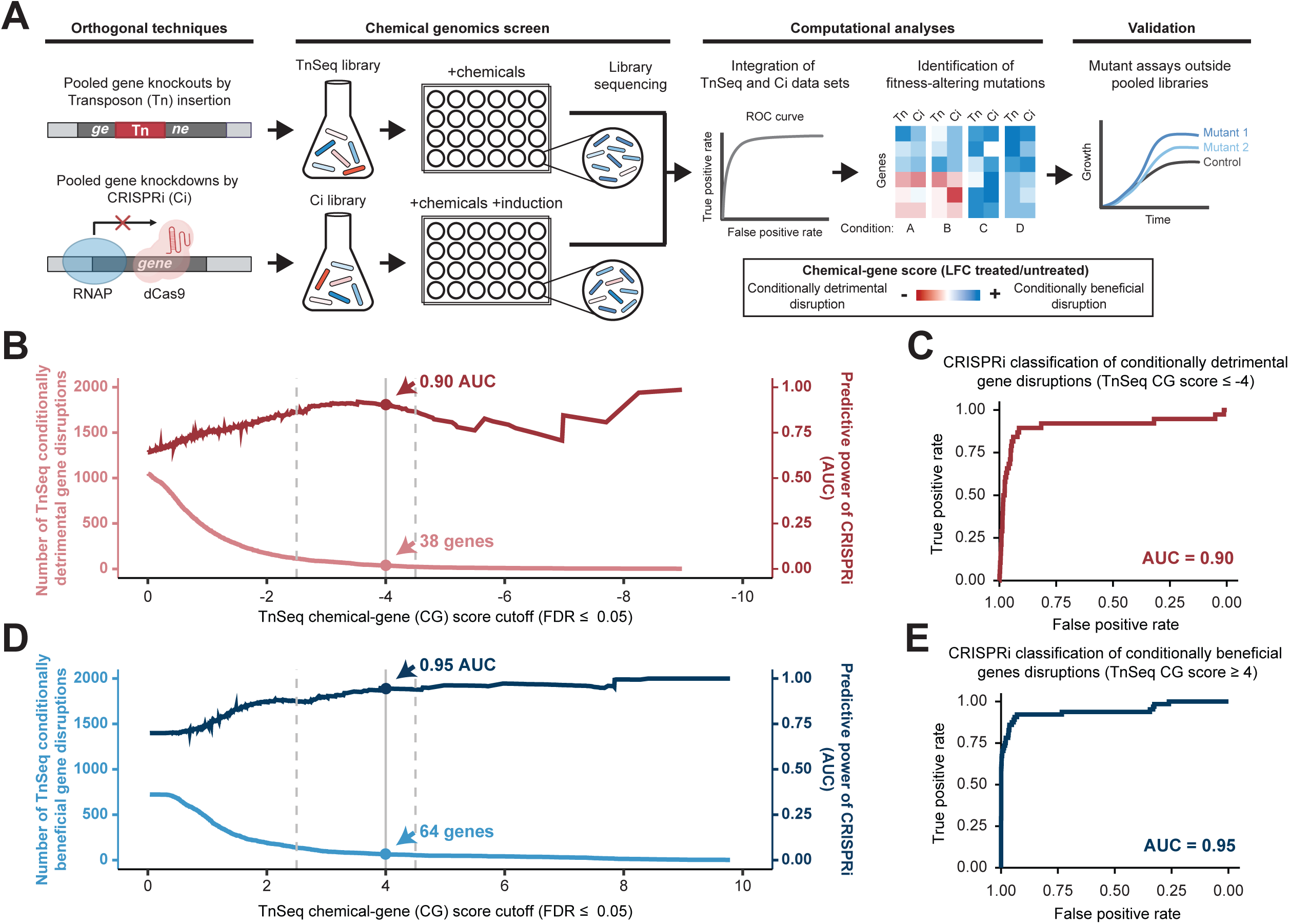
Parallel Z. mobilis chemical genomics screens using orthogonal TnSeq and CRISPRi libraries. **A)** Schematic of integrative chemical genomics using TnSeq and CRISPRi libraries grown with fermentation-relevant chemicals. RNAP, RNA polymerase. **B and D)** ROC curve area under the curve (AUC) across chemical-gene (CG) score cutoff values for conditionally detrimental (B) and conditionally beneficial (D) gene disruptions, using the TnSeq library as ground truth. The darker line depicts AUC at a given CG score cutoff (right y-axis) while the lighter line represents the number of genes which pass the associated cutoff value (left y-axis). Dashed vertical lines approximate the bounds of a flexible range within which researchers may choose to select cutoffs by weighing the tradeoff between number of hits and reliability of those hits. The solid gray line at |CG score| = 4 marks the score cutoff chosen in this work to identify genes for further study. **C and E)** ROC curves generated using (C) CG score ≤ −4 and (E) CG score ≥ 4 using the TnSeq library as ground truth.

Next, we systematically determined a CG score cutoff to apply across libraries and reliably define engineering targets. To do so, we constructed receiver operating characteristic (ROC) curves that test the ability of one library to accurately predict results from the other, where higher area under the curve (AUC) represents greater prediction accuracy. This strategy was previously used to evaluate the ability of an *Escherichia coli* CRISPRi library to classify known essential genes based on gold-standard deletion libraries (31). For each of our libraries, we designated gene disruptions as either conditionally detrimental (disruption decreases fitness; negative CG scores) (Fig. 1B-C) or conditionally beneficial (disruption increases fitness; positive CG scores) (Fig. 1D-E) along a range of CG score cutoffs (and FDR ≤ 0.05) and calculated AUC for ROC curves at each cutoff. As the cutoff increases, the number of genes passing a given cutoff decreases while predictive power generally increases until the gene sample size becomes insufficient, causing noise in the AUC calculation (e.g., Fig. 1B). Therefore, there is a tradeoff between the number of genes identified and the reliable classification of gene phenotypes, and researchers may opt for more stringent or relaxed cutoffs depending on their tolerance for false results and bandwidth for follow-up experiments. For this study, we chose a cutoff of |CG score| ≥ 4. At this cutoff, CRISPRi library CG scores reliably classify TnSeq mutant phenotypes (AUC = 0.90 for 38 conditionally detrimental gene disruptions and AUC = 0.95 for 64 conditionally beneficial gene disruptions) (Fig. 1B, Fig. 1D, Fig. S3). In the same manner, TnSeq CG scores are able to reliably classify CRISPRi mutant phenotypes at this cutoff (AUC = 0.86 for 43 conditionally detrimental gene disruptions and AUC = 0.94 for 65 conditionally beneficial gene disruptions) (Fig. S3). Applying this cutoff, we identified 103 genes with significant CG scores (|CG score| ≥ 4, FDR ≤ 0.05) in at least one chemical condition and genetic library (Fig S3, Tables S2 and S3). Of the 103 significant genes, 31 had significant CG scores in both libraries for at least one condition (Fig 2A). We consider these genes with cross-validated phenotypes to be potential engineering targets for generating inhibitor-resistant production strains. The other 72 genes were significant in one library but not in the other and may warrant further investigation (Table S3 and Fig. S4).

**Figure 2.**
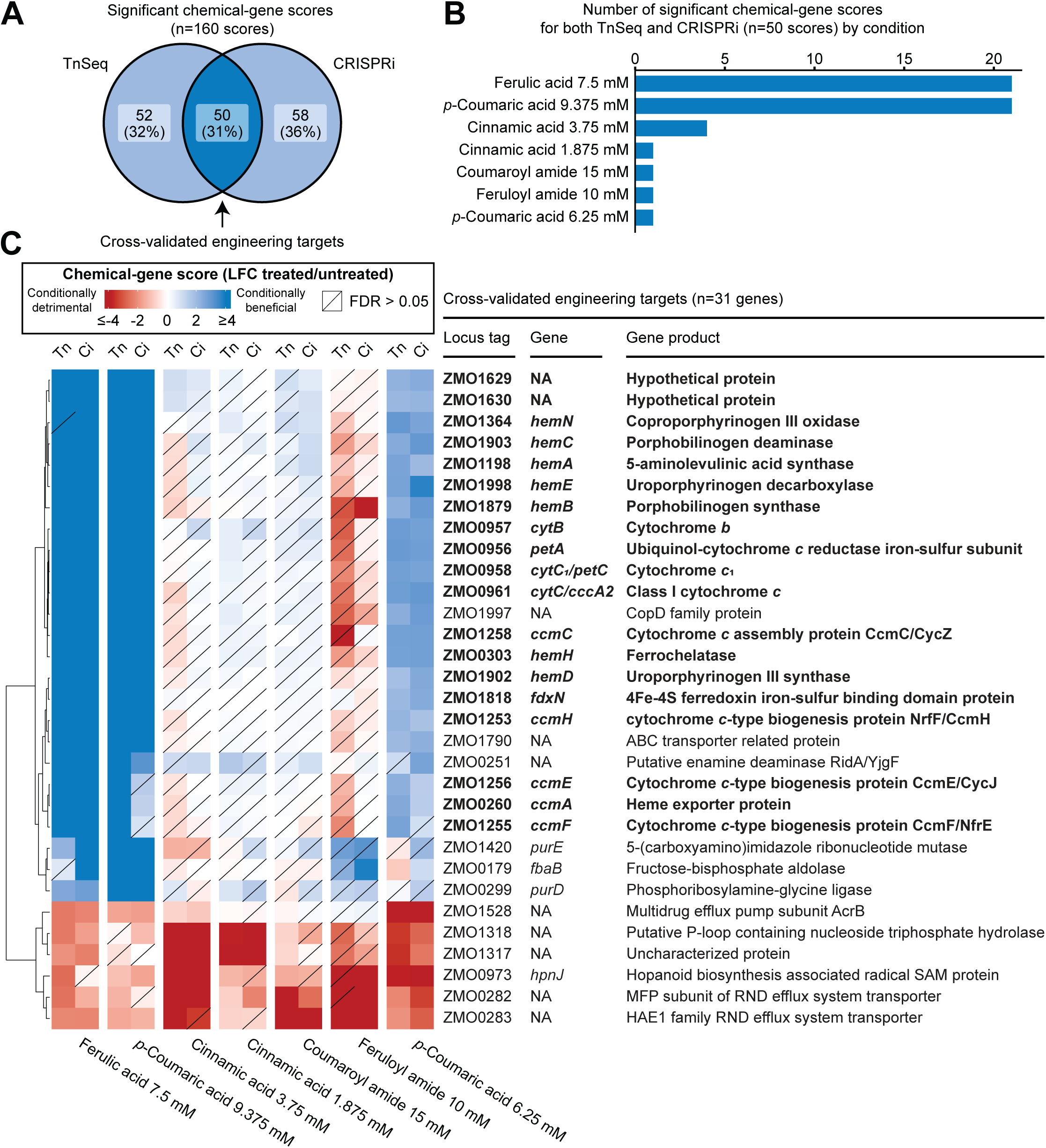
Significant chemical genomics hits identified by TnSeq and CRISPRi libraries. **A)** Venn diagram showing number of significant chemical gene scores identified by either TnSeq or CRISPRi, or in both. **B)** Chart depicting number of significant genes (|chemical-gene score| ≥ 4 in both TnSeq and CRISPRi) in each condition that had at least one significant gene. **C)** Hierarchical clustered heatmap showing chemical-gene scores for significant genes across conditions that had at least one significant gene. Bolded genes are also displayed in Fig. 3A. Tn, TnSeq library; Ci, CRISPRi library. Diagonal lines indicate statistical non-significance (FDR > 0.05). Hierarchical clustering was applied only to the y-axis (genes axis). Conditions are arranged from left to right by the number of significant genes, as displayed in (A).

### Disruption of the cyt *bc*_1_/cyt *c* electron transport pathway increases fitness in phenolic acids

Of the 31 cross-validated engineering targets, 21 gene disruptions improved fitness in ferulic acid and *p*-coumaric acid (Fig. 2B-C, Fig. S5). Strikingly, these 21 genes include all known structural genes needed to synthesize a quinol-oxidizing electron transport pathway consisting of a cytoplasmic membrane-bound cytochrome *bc_1_* complex (*ZMO0956-ZMO958*) and a periplasmic cytochrome *c* (*ZMO0961*) (herein termed the cyt *bc_1_*/cyt *c* pathway). Notably, what electron transfer components this pathway interacts with and ultimately any final electron acceptors are unknown. Other genes required for maturation of these respiratory enzymes had similarly positive CG scores in these conditions, including genes encoding synthesis of heme, an essential cofactor for cytochrome-mediated electron transfer (32–34), and the cytochrome *c* maturation (*ccm*) complex, required for covalent heme attachment to cytochromes *c* and translocation to the periplasm (35–37). Of the remaining genes in this group, hierarchical clustering of CG scores across all conditions revealed that the cytoplasmic ferredoxin *fdxN* (*ZMO1818*) and two members (*ZMO1629, ZMO1630*) of an operon of unknown function (*ZMO1631-ZMO1628*) clustered with cyt *bc_1_*/cyt *c* pathway genes (Fig. S4). While *ZMO1631* is annotated as a TonB-dependent receptor, its downstream genes (*ZMO1630-ZMO1628*) are poorly annotated. ZMO1630-1628 are predicted to have α-helices indicative of localization to the cytoplasmic membrane. Furthermore, AlphaFold3 modeling (38) predicts that ZMO1630-ZMO1628 form a transmembrane complex (ipTM = 0.86, Fig. S6), but are not predicted to associate with ZMO1631 (ipTM = 0.57). These uncharacterized genes were included in follow-up analyses due to their similar phenotypes, which suggest they have related functions. Importantly, genes encoding other quinol-oxidizing pathways, such as cytochrome *bd* oxidase (*ZMO1571-1572*) and cytochrome *c* peroxidase (*ZMO1136*), did not show significant phenotypes in any condition, indicating that the increased phenolic acid resistance phenotype is specific to the cyt *bc_1_*/cyt *c* pathway rather than electron transport pathways in general (Fig. S7). These fitness patterns were specific to phenolic acids and did not extend to non-phenolic cinnamic acid or the phenolic amides, feruloyl and coumaroyl amide (Fig 2C).

Iron-sulfur clusters also play an indispensable role in the cyt *bc_1_*/cyt *c* pathway, with the Rieske iron-sulfur protein (*ZMO0956*) and *hemN* utilizing [2Fe-2S] and [4Fe-4S] clusters, respectively (39,40). Additionally, the ferredoxin *fdxN* utilizes a [4Fe-4S] cluster (41). We therefore reasoned that disruption of Fe-S cluster biosynthesis should provide a similar fitness benefit with phenolic acids. However, genes encoding the iron-sulfur cluster biosynthetic Suf system (42) are essential and therefore are absent from our TnSeq and CRISPRi library comparisons since Tn insertion into essential *suf* genes is lethal (43 and this study). The notable exception to this is the first gene in the *suf* operon, *ZMO0422*, which encodes a non-essential predicted transcription factor (Fig. S8). While TnSeq mutants of *ZMO0422* showed minimal phenotypic effects, CRISPRi mutants exhibited strong positive scores in the presence of ferulic and *p*-coumaric acids, likely due to reduced transcription of the downstream *suf* genes. Consistent with this, partial knockdown of *suf* genes using mismatched sgRNAs in the CRISPRi library resulted in similarly positive phenotypes with ferulic and *p*-coumaric acids (Fig. 3A), confirming that hindering Fe-S cluster biosynthesis also increases fitness with these compounds, possibly by disrupting maturation of the cyt *bc*_1_/cyt *c* pathway.

**Figure 3.**
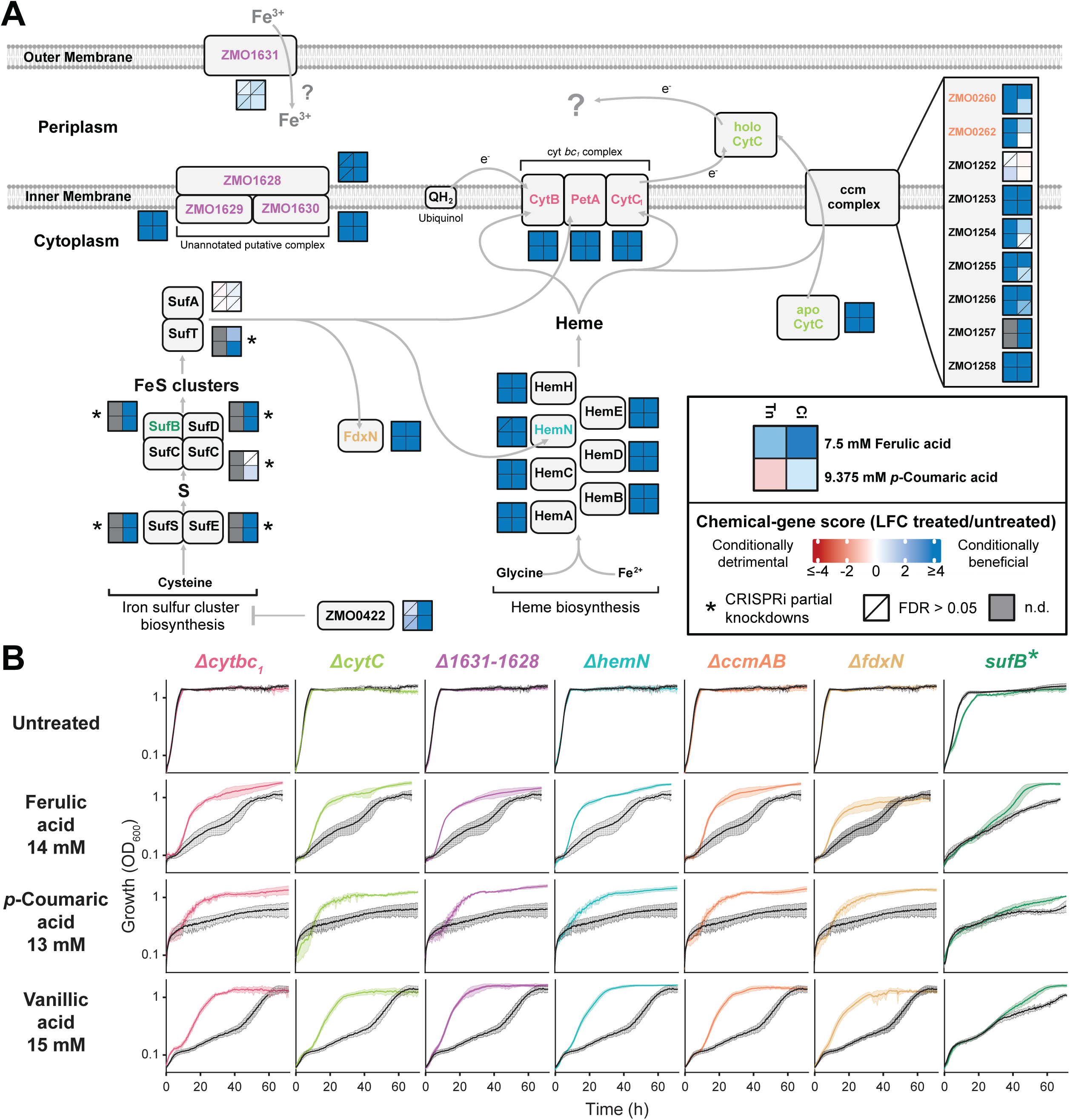
Fitness and growth of functionally related gene mutants in ferulic and *p*-coumaric acid. **A)** Pathway map showing functional relation between genes which had significant phenotypes in 7.5 mM ferulic and 9.375 mM *p*-coumaric acid. Heatmaps next to each gene depict chemical-gene scores in both conditions for TnSeq and CRISPRi libraries. Tn, TnSeq library; Ci, CRISPRi library; n.d., no data. Diagonal lines indicate statistical non-significance (FDR > 0.05). Heatmaps with asterisks display the median CG score for CRISPRi partial knockdown mutants using mismatched sgRNAs. n.d. denotes no data. Corresponding locus tags for protein names shown are: SufB, ZMO0423; SufC, ZMO0425; SufD, ZMO0426; SufS, ZMO0427; SufT, ZMO0428; SufA, ZMO0429; SufE, ZMO1067; HemA, ZMO1198; HemB, ZMO1879; HemC, ZMO1903; HemD, ZMO1902; HemN, ZMO1364; HemE, ZMO1998; HemH, ZMO0303; CytB, ZMO0957; PetA, ZMO0956; CytC1, ZMO0958; CytC, ZMO0961; FdxN, ZMO1818. **B)** Growth curves displaying anaerobic growth with phenolic acids for deletion mutants or a sufB Mismatch-CRISPRi partial knockdown mutant (colored lines) and wild type *Z. mobilis* or a non-targeting CRISPRi control strain, respectively (black lines). Growth curve error bars are standard deviation of quadruplicate samples.

We next validated that disruption of the cyt *bc_1_*/cyt *c* pathway improves fitness against phenolic acids in a non-competitive assay. We first constructed in-frame deletion null mutants for a subset of genes encoding the pathway and its maturation via homologous recombination (Table S1) and compared the growth of these deletion mutants to a wild type (WT) parent when challenged with ferulic or *p*-coumaric acids under anaerobic conditions (Fig 3B). We also constructed and tested two CRISPRi partial knockdown strains of the *suf* operon using mismatched sgRNAs targeting *sufB*, the first biosynthetic gene downstream of *ZMO0422* (Fig 3B and Fig S9). Finally, to interrogate whether the mutations increased fitness to other phenolic acids, we also measured growth with vanillic acid. In untreated media, all deletion mutants grew similarly to WT *Z. mobilis*, demonstrating that loss of these genes had no fitness cost during anaerobic growth in rich medium. As expected for essential gene perturbations, *suf* partial knockdown strains showed slight growth defects in untreated media compared to a non-targeting CRISPRi control strain. However, in the presence of any of the phenolic acids, including vanillic acid, all mutants grew more rapidly and/or to a higher terminal optical cell density (OD_600_) than their parent control (Fig. 3B).

### Ferulic acid fails to induce a strong iron starvation response in *Z. mobilis*

There is mixed evidence on whether phenolic acids can chelate iron, which could reduce iron bioavailability and perturb iron homeostasis (44,45). Given that our target genes were iron-containing cellular components, we hypothesized that the improved fitness of our deletion and knockdown mutants in phenolic acids could result from sparing iron for other essential processes. To test the ability of ferulic, *p*-coumaric and vanillic acids to chelate iron, we performed a chrome azurol S (CAS) iron binding assay (46). This assay showed negligible iron binding at concentrations approximating those used in our chemical genomics screen and mutant growth experiments (9.375-18.75 mM), except for a slight iron binding signal for 18.75 mM vanillic acid (Fig. 4A-B and Fig. S10). To examine possible iron stress directly in *Z. mobilis*, we measured expression of a *lacZ* fusion to the iron-starvation responsive *fhuE (ZMO0188*) promoter (Isabel Askenasy and Patricia J. Kiley, manuscript in preparation); however, we observed only a minor increase in promoter activity with 10mM ferulic acid, a concentration sufficient to slow growth and ∼670x higher than concentration of the known iron chelator, 2,2’ dipyridyl, which efficiently induced *fhuE* expression (Fig. 4C). Overall, our results suggest that phenolic acids have a very minor effect on iron bioavailability.

**Figure 4.**
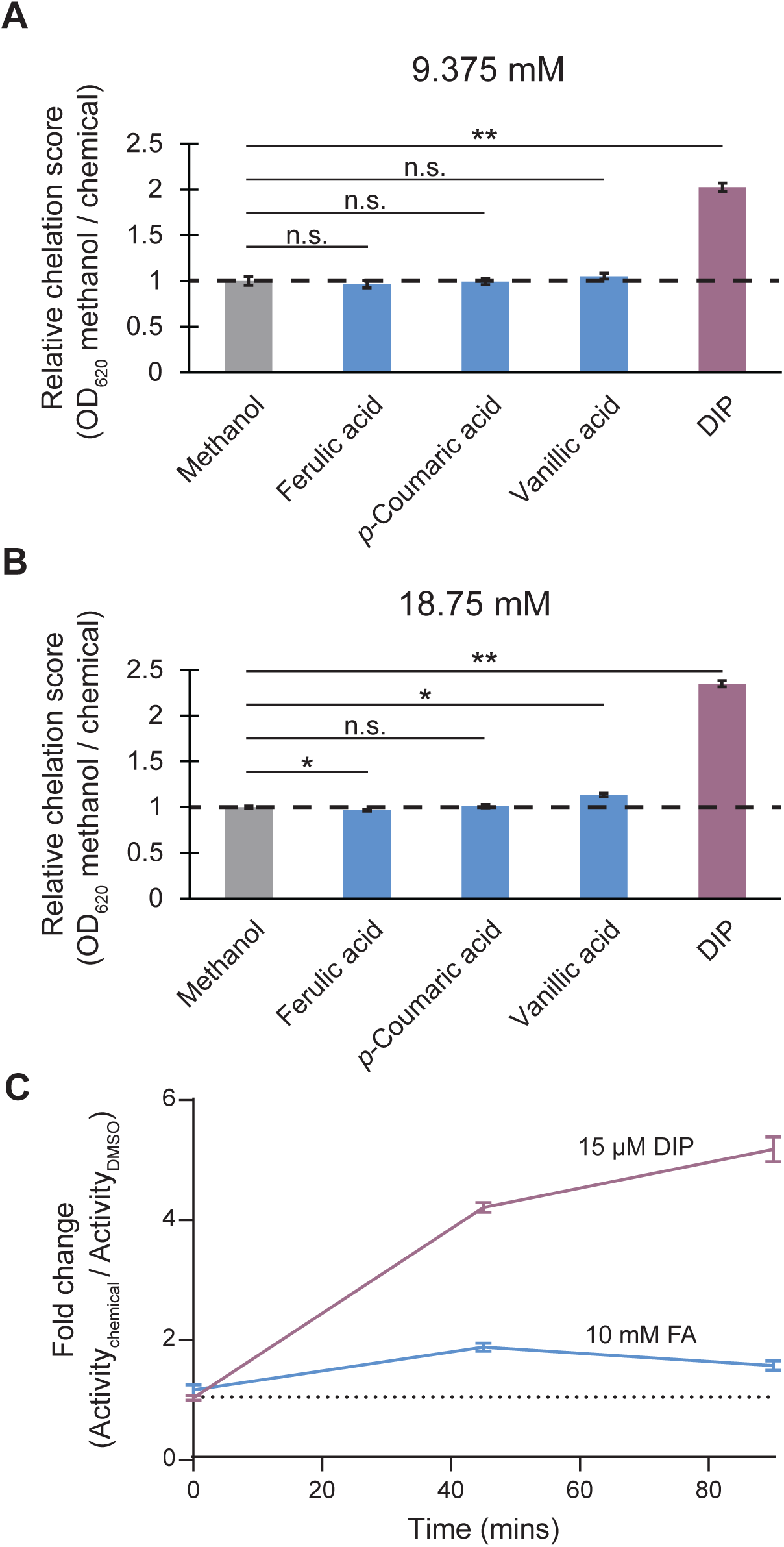
Phenolic acid iron chelation assays. **A and B)** Results of an in vitro colorimetric chrome azurol S (CAS) iron binding assay with chemicals at (A) 9.375 mM or (B) 18.75 mM, concentrations approximating those used in the chemical genomics screen and growth experiments (blue). The known metal chelator 2,2’ dipyridyl (DIP) was used as a positive control (purple). Methanol was used as a solvent control (gray). Dashed line indicates the average of the solvent control. **C)** β-galactosidase assay measuring *fhuE* promoter activity with ferulic acid or DIP. Activity is shown as fold change in Miller units between a 1% DMSO negative control and treatment. Error bars are representative of standard error of triplicate samples.

### Ferulic acid causes proteome remodeling in the cell envelope

To further investigate the cellular effects of phenolic acid induced stress, we performed untargeted proteomics and compared protein abundance between untreated and ferulic acid-treated WT, Δ*cytbc_1_* (*ΔZMO0956-ZMO0958*) and Δ*ZMO1631-ZMO1628*, the operon encoding proteins of unknown function. The three strains were grown anaerobically and treated with a sub-lethal concentration of ferulic acid. Samples for proteomics were collected just prior to treatment and two hours afterwards. In total, our proteomic analysis detected peptides corresponding to the vast majority of predicted ORFs in *Z. mobilis* (1,622 proteins detected of the 1,852 predicted ORFs, Fig. 5A). We compared protein relative abundance between samples treated with ferulic acid/DMSO or DMSO after two hours, revealing 57 proteins that had significantly changed abundance in response to ferulic acid in WT *Z. mobilis* (FDR < 0.05, |log_2_ fold change| > 1). The Δ*cytbc_1_* and Δ*ZMO1631-ZMO1628* mutants showed 42 and 75 significantly changed proteins, respectively (Fig. 5B).

**Figure 5.**
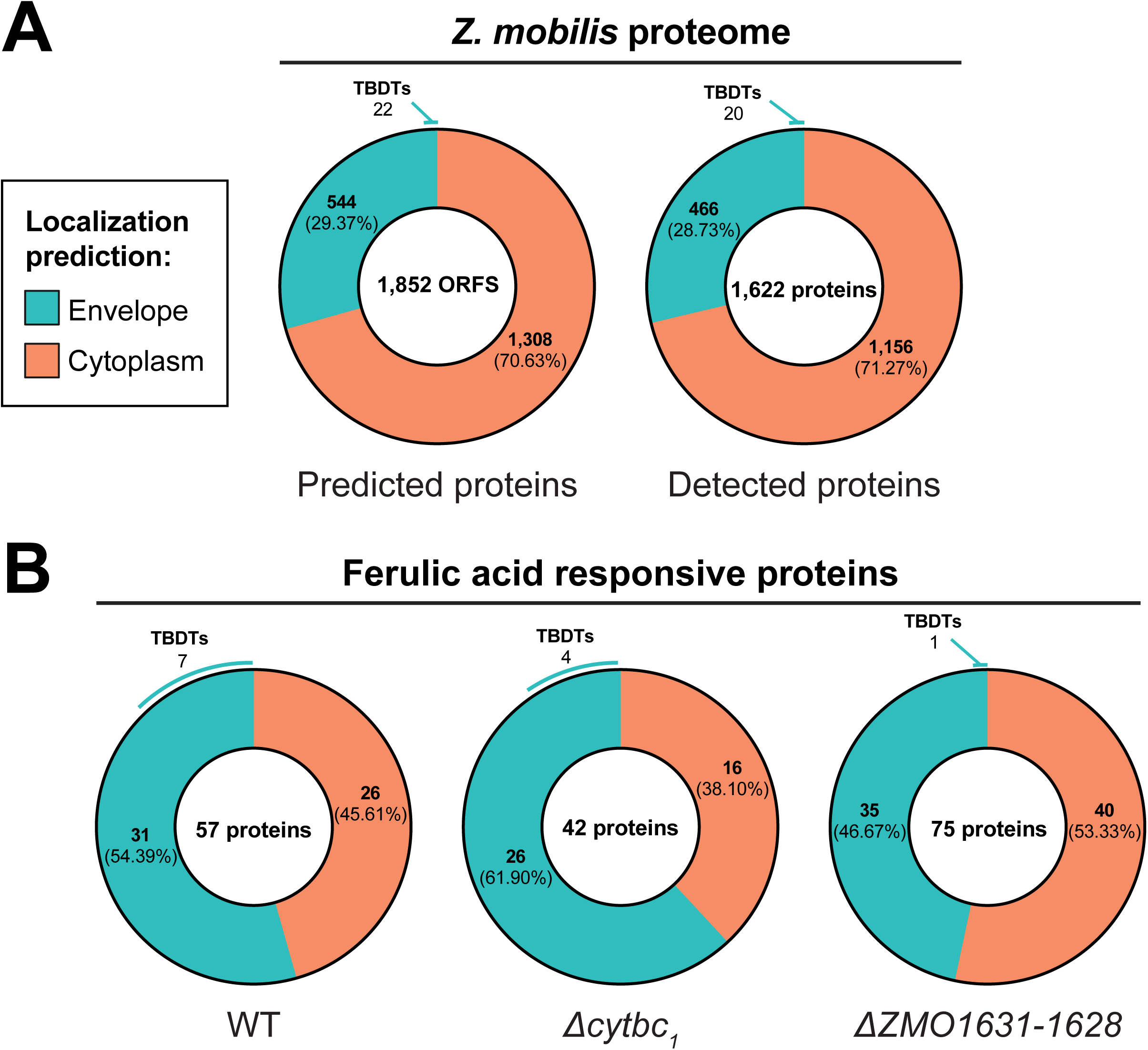
Proteome profiling of ferulic acid-treated *Z. mobilis.* **A)** Protein localization prediction for the putative *Z. mobilis* proteome (left) and for proteins which were detected in at least one sample in the proteomics screen (right). **B)** Protein localization prediction for proteins which had significant changes in abundance (FDR < 0.05, |LFC| > 1) in response to ferulic acid treatment in WT, Δ*cytbc1*, and Δ*ZMO1631-ZMO1628.* Number of TonB-dependent transporters (TBDTs) which had significantly changed abundance are also shown.

The overall proteomic response to ferulic acid was similar across the three strains tested, with many ferulic acid-responsive proteins predicted to be localized to the cell envelope (inner membrane, periplasm, outer membrane). We used SignalP 6.0 and DeepTMHMM 1.0 (47,48) to predict the localization of all *Z. mobilis* proteins. Proteins containing a secretion signal sequence, e.g., Sec or Tat peptides that target proteins for secretion to the envelope (49), or a transmembrane domain consisting of either α-helical or β-barrel domains were annotated as envelope proteins. All other proteins were annotated as cytoplasmic. Our analysis found that 544 of the predicted 1,852 ORFs (29.37%) in *Z. mobilis* encode proteins predicted to localize to the cell envelope (Fig. 5A, Table S4). Of the 1,622 proteins that were detected in all proteomics samples, 466 (28.73%) are predicted envelope proteins. Notably, envelope proteins made up a disproportionately large fraction of ferulic acid responsive proteins across all three strains (54.39% in WT, 61.90% in *Δcytbc_1_*, and 54.39% in *ΔZMO1631-ZMO1628*) (Fig. 5B). These percentages are significantly higher than would be expected by random chance (P = 0.0074, 0.0041, 0.0431 for WT, *Δcytbc_1_* and *ΔZMO1631-ZMO1628,* respectively, Fischer’s exact test, Fig S11), suggesting that ferulic acid stress targets the cell envelope.

### Transporter abundance is modulated in response to treatment with ferulic acid

To further identify trends in proteome remodeling in response to ferulic acid, we performed functional enrichment analysis using the STRING database (50) (Fig. S12, Table S5). Functional enrichments were similar across all three strains, indicating that the mutants respond to ferulic acid similarly to WT despite their increased phenolic acid resistance. These common enrichments include cell envelope proteins, oxidoreductase activity and carbon metabolism. In particular, TonB-dependent transporters (TBDTs) were highly represented among the functional enrichments, suggesting a key role in responding to ferulic acid stress. TBDRs are proton motive force (PMF) coupled complexes that facilitate transport of large compounds such as siderophores, vitamin B12 and glycans across the outer membrane (51–53). Of the 22 annotated TBDTs in *Z. mobilis*, seven had significantly decreased abundance in at least one strain in response to ferulic acid (|log_2_ fold change| > 1, FDR < 0.05, Fig. 6).

**Figure 6.**
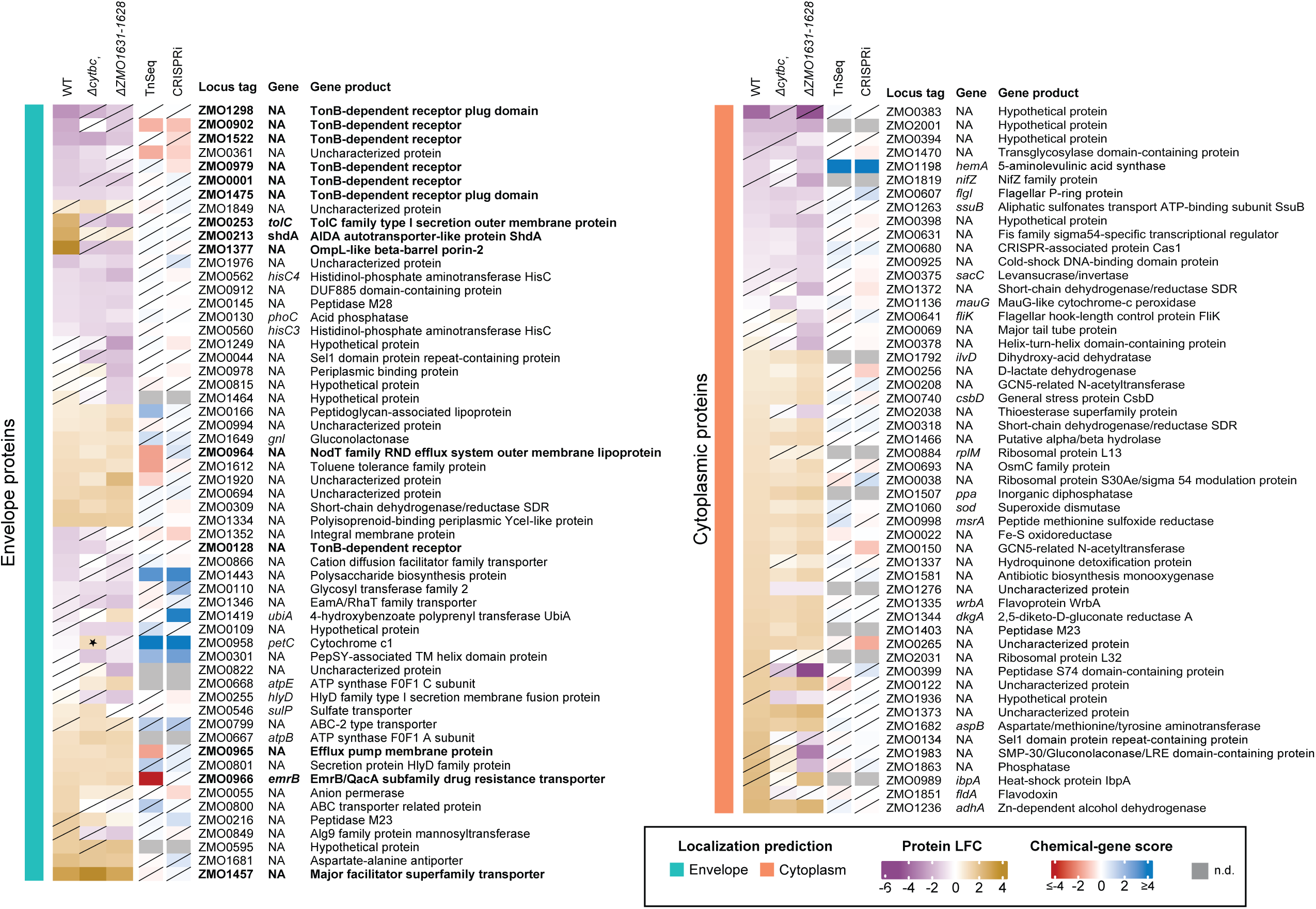
Heatmap of ferulic acid-responsive proteins. Heatmap of all genes which had significantly (FDR > 0.05, |LFC|>1) changed abundance in at least one strain in response to ferulic acid treatment. Chemical-gene scores in 7.5 mM ferulic acid are displayed in the two right-most columns. n.d., no data. Transporter proteins that are discussed in the main text are bolded. The starred protein is deleted in the associated strain, and yet has a significant LFC due to missing value imputation during proteomic analysis.

Our proteomics data also revealed several transporters that increased abundance across all three strains in response to ferulic acid, including a predicted major facilitator superfamily transporter (ZMO1457) and a predicted efflux system (ZMO0964-ZMO0966) (Fig. 6, Table S6). Other transporters were responsive only in a single strain. For example, ZMO0253, ZMO0213 and ZMO1377 are transport-related proteins that increase abundance in WT in response to ferulic acid but were not responsive to ferulic acid in *Δcytbc_1_* or *ΔZMO1631-ZMO1628*. While transporters that increase in abundance in response to ferulic acid might be candidates for proteins involved in efflux of the chemical, none of the genes encoding these transporters showed consistently strong CG scores across the Tn and CRISPRi genetic screens (Fig. 6). This could be explained by possible functional redundancy among multiple transporters, though further study is required to genetically dissect this.

## DISCUSSION

Industrial microorganisms resilient to plant- and process-derived chemical stressors are required for efficient bioproduction. However, incomplete understanding of the genetic basis for stress tolerance and susceptibility hinders rational engineering of robust strains. To identify genetic engineering targets, we developed an orthogonal and parallel screening strategy using TnSeq and CRISPRi libraries that leverages the complementary strengths of both technologies to identify target genes with high specificity. By applying our strategy to investigate the effects of inhibitory chemicals on *Zymomonas mobilis* during anaerobic growth, we narrowed the genetic search space from 103 genes identified by at least one library to 31 cross-validated engineering targets. All follow up mutants displayed the predicted chemical resilience phenotypes, indicating that our strategy precisely identifies true positives. Our combined approach is readily applicable to a wide range of bacteria with implications across industrial, medical, and environmental research—especially where reducing frequency of false hits is highly desirable—and could be immediately implemented for species with existing CRISPRi and TnSeq libraries, such as *E. coli, Bacillus subtilis* or *Streptococcus pneumoniae* (31, 54–58).

This work builds on prior investigations by characterizing chemically mediated stress in production-relevant anaerobic conditions. Previously, a transposon screen conducted aerobically identified gene disruptions that decrease fitness in common feedstock inhibitors, including ferulic and *p*-coumaric acids (15). Among these are genes encoding cytochrome *c* maturation (*ccmF, ZMO1255; ccmI, ZMO1252*; *ccmH, ZMO1253*) and Fe-S cluster biosynthesis (*sufE, ZMO1067; sufA, ZMO0429*). In contrast, we found that disrupting these genes improved fitness with ferulic and *p*-coumaric acids under anaerobic conditions. Given that *Z. mobilis* displays distinct physiology depending on oxygen and nutrient availability (15, 43, 59, 60), this contrast is likely due to differences in growth conditions (e.g., aerobic versus anaerobic growth and media composition). Our identification of more conditionally beneficial gene disruptions than Skerker and Leon et al. may also reflect our use of contemporary quantification technologies—next generation sequencing instead of microarrays—which improved our ability to measure mutant abundance.

Our dual-library approach revealed that disrupting the cyt *bc*_1_/cyt *c* electron transport pathway improves fitness of *Z. mobilis* in phenolic acids under anaerobic conditions, indicating an unexpected anaerobic function for this pathway. In mitochondria and many bacteria, this pathway is well characterized for its role in aerobic respiration, where it transfers electrons to cytochrome *c* oxidase. However, in *Z. mobilis*, this pathway likely does not contribute directly to aerobic respiration, as the organism lacks a cytochrome *c* oxidase and instead uses a ubiquinol-cytochrome *bd* oxidase to reduce O_2_ (61). An anaerobic function for the cyt *bc_1_*/cyt *c* pathway has been established in a few bacteria that carry out anaerobic respiration. For example, in *Pseudomonas aeruginosa,* electrons can be transferred from the cyt *bc_1_*/cyt *c* pathway to nitrite, nitric oxide, or nitrous oxide via associated reductases during anaerobic growth (62). However, no equivalent reductases are annotated in *Z. mobilis*, nor did our screen reveal any terminal reductase candidates, leaving the anaerobic function of this pathway an area for further investigation.

Our study also identified an operon of unknown function (*ZMO1631-ZMO1628*) with anaerobic phenotypes similar to cyt *bc_1_*/cyt *c* pathway phenotypes, suggesting a functional relationship. *ZMO1629* and *ZMO1630* had significant chemical-gene scores in both Tn and Ci libraries, and deletion of the entire operon confers increased fitness to ferulic and *p*-coumaric acids. The operon is predicted to be regulated by the iron starvation-responsive transcription factor, Fur (63), suggesting that it could play a role in iron acquisition and retention. Indeed, *ZMO1631* is a predicted TonB-dependent receptor, whose function is often associated with siderophore-mediated iron transport. AlphaFold3 modeling predicts that *ZMO1630-1628* form a 12-transmembrane helix complex, with *ZMO1628* also containing a predicted PepSY-domain, suggesting a possible function in transport. Moreover, other iron regulated PepSY-domain containing proteins from *P. aeruginosa* and *Bradyrhizobium japonicum* have been identified as cytoplasmic membrane-embedded ferric reductases which are involved in siderophore-mediated iron uptake (64–66). Thus, it is possible that this gene cluster may have a similar function in iron acquisition in *Z. mobilis*.

We further characterized the phenolic acid stress response using proteomics and found that ferulic acid causes remodeling of the cell envelope proteome, which is consistent with previous literature reporting that phenolic acids can perturb cell membranes (5, 67, 68). Furthermore, ferulic acid treatment results in changes in abundance of several transporters, including putative efflux systems and TonB-dependent transporters, suggesting a potential role in ferulic acid transport. In the closely related genus *Sphingobium*, TonB was shown to play a role in uptake of ferulic acid and other lignin-derived phenolics for subsequent catabolism (69, 70). However, specific TBDRs involved in ferulic acid uptake were not identified. *Z. mobilis* is not known to catabolize phenolic acids, and while *tonB* (*ZMO1717*) was essential in our TnSeq data—prohibiting further phenotypic analysis in our comparative screen—CRISPRi library partial knockdowns of *tonB* with mismatched sgRNAs had statistically insignificant ferulic acid phenotypes (Table S2). Consequently, it is possible that if TBDTs are involved in phenolic acid uptake in *Z. mobilis*, it is as an unintended substrate. Interestingly, the *ΔZMO1631-ZMO1628* mutant, which harbors a deletion for the annotated TBDT gene *ZMO1631*, had fewer significantly downregulated TBDTs than the WT or *Δcytbc_1_* strains. It is possible that this deletion might also provide resistance by decreasing chemical import. Although our genetic screens did not pinpoint other TBDTs with phenolic acid resistance phenotypes, this could be explained by redundant functions among multiple TBDTs.

Our orthogonal screening approach robustly identified genes that provided accurate engineering targets. However, genes that fell below our cutoff and were identified only by CRISPRi or TnSeq may also be beneficial engineering targets. Thus, if a more comprehensive view of all potential targets is desired, then adjusting the cutoffs to be more permissive is straightforward but will need to be counterbalanced by screening more false positives and utilizing more resources. CRISPRi and TnSeq are known to each have certain limitations in associating phenotypes with particular genes, making the dual approach used here so effective in minimizing false positives. In the case of CRISPRi, off-target sgRNA binding or toxic sgRNA sequences (71) can produce incorrect phenotypes. For Tn insertions, multiple insertions per genome (72) or stabilization of merodiploid/polyploid states of the chromosome caused by insertions in essential genes (15, 73, 74) can also obscure phenotype assignments. Finally, differing *cis* effects on downstream gene expression between the two techniques may also yield different gene phenotypes revealing interesting biology. For example, while Tn insertion in *ZMO0422*—the first gene in the *suf* operon and the only one not directly performing iron sulfur cluster assembly—had an insignificant phenotype with phenolic acids, *ZMO0422* CRISPRi knockdowns were conditionally beneficial. Given that our Mismatch-CRISPRi partial knockdowns of essential *suf* genes also provided a conditional relative fitness advantage, the CRISPRi phenotype for *ZMO0422* is likely due to CRISPRi polarity onto downstream *suf* genes.

In conclusion, we developed an approach for rapid identification of high-value gene targets and used it to advance our understanding of *Z. mobilis* chemical tolerance in anaerobic conditions, providing valuable insights for future bioengineering efforts. We highlight the cyt *bc*_1_/cyt *c* pathway as a promising target for engineering robust biofuel production strains with improved phenolic acid resistance, particularly in plant-based feedstocks with high concentrations of phenolics (4, 75, 76). Looking ahead, we anticipate that adaptations of our comparative approach will accelerate discoveries in fields driven by chemical-gene interactions, including industrial biomanufacturing, combating antibiotic resistance, and understanding chemically-mediated symbioses within human or plant microbiomes.

## MATERIALS AND METHODS

### Bacterial growth conditions

*Z. mobilis* was grown at 30°C in either *Zymomonas* Rich Defined Medium (ZRDM) or Rich Medium Glucose (ZRMG) (43). Anaerobic growth was achieved either by culturing in a Coy chamber with 10% CO_2_, 5% H_2_, and balance N_2_ or by sparging with 95% N_2_ and 5% CO_2_*. Escherichia coli* was grown in Lennox broth at 37°C. Where indicated, a final concentration of 100 µg/mL ampicillin (amp) or 20 µg/mL chloramphenicol (cm) was added to *E. coli* media, and 100 µg/mL cm to *Z. mobilis* media. A final concentration of 300 µM diaminopimelic acid (DAP) was added for growth of Dap^−^ *E. coli* mating strains. Where indicated, a final concentration of 1 mM isopropyl β-D-1-thiogalactopyranoside (IPTG) was included.

For anaerobic growth measurements in a Coy chamber, 24-well plates containing 1.2 mL ZRDM supplemented with 1% v/v DMSO or phenolic acid dissolved in DMSO (100x) were inoculated after equilibration. Strains were grown to saturation, and plates were inoculated with saturated cultures to a final O.D._600_ of 0.05. Growth of strains was analyzed in quadruplicate for each condition by measuring O.D._600_ of each well every thirty minutes for 72 hours using a Tecan Spark plate reader equipped with a stacker.

### Construction of bacterial strains and libraries

Strains are listed in Table S1. Deletion mutants were constructed via homologous recombination by screening for the loss of either GFP (26) or selecting for the loss of *sacB* (manuscript in preparation). Deletion mutants were verified by whole genome resequencing using NextSeq1000 (77).

Individual Mobile-CRISPRi mutants were constructed as described previously (28, 78). Briefly, sgRNA-encoding sequences were cloned between BsaI sites of the Mobile-CRISPRi vector (pJMP2656), and the Mobile-CRISPRi system was transferred to Tn*7att* site of the *Z. mobilis* chromosome via conjugation.

Construction of a *Z. mobilis* Tn5-based insertion library for use in TnSeq analysis was carried out with ZRMG under aerobic conditions as previously described (79). Construction of the *Z. mobilis* Mobile-CRISPRi library was described previously and consists of three sub-libraries: (1) sJMP2618, Z1-genes, perfect-match spacers; (2) sJMP2619, Z2-controls, non-targeting spacers; and (3) sJMP2620, Z3-mismatches, mismatched spacers (43).

### Chemical genomics screen

Chemical genomics screens were performed anaerobically, and trace O_2_ was removed by placing all components inside the Coy anaerobic chamber for at least 18 hours before inoculation. Relevant concentrations of chemicals and suppliers are listed in Table S1. Feruloyl amide and coumaroyl amide were synthesized as previously described (4). Chemical concentrations that caused ∼10-40% reduction in the empirical area under the growth curve, as calculated by the Growthcurver R package (v0.3.1), for *Z. mobilis* sJMP2554 (non-targeting CRISPRi strain) compared to the untreated control were employed in the screen (80).

The *Z. mobilis* CRISPRi library (sJMP2618, 2619, 2620) inoculum was prepared as follows: sub-libraries Z1, Z2 and Z3 were mixed at a ratio of 8:1:6 and 400 µL total library was added to 40 mL ZRDM and grown at 30°C with stirring to O.D._600_ ∼0.5. Untreated control samples were collected for sequencing, and cells from this flask were used to inoculate 1.5 mL ZRDM with 1 mM IPTG in the presence or absence of chemical to O.D._600_ ∼0.01 in 24-well plates and grown in a Tecan Spark multimode microplate reader with Spark Stacker, shaking for 10 seconds every 30 minutes, to stationary phase (∼46 hours). To allow sufficient generations for depletion of gene product following induction of the CRISPRi system with IPTG, the process was repeated by sub-culturing into a second 24-well plate and growing from a starting O.D._600_ ∼0.01 for another ∼46 hours. At this time, CRISPRi library samples with ∼8-30% reduction in the empirical area under the growth curve (80) compared to the untreated controls were harvested for sequencing by centrifuging and storing cell pellets at −20°C.

The *Z. mobilis* Tn5 transposon library (PK15455) was screened similarly, except that no IPTG was added to growth media, and 100 µL of library was initially diluted in 10mL ZRDM to O.D._600_ ∼1.8 before transferring 75 µL to each 24-well plate for a starting O.D._600_ ∼0.09. Growth and subculturing into the second 24-well plate was performed as was done for the CRISPRi library. Samples were collected for sequencing at the same time points as CRISPRi samples by centrifuging and storing cell pellets at −20°C.

### TnSeq library sequencing and data analysis

TnSeq library genomic DNA was extracted using Qiagen DNeasy Blood and Tissue Kit (Qiagen, 69504), fragmented using a Covaris S220 Focused-Ultrasonicator, converted to blunt-end DNA using NEBNext End Repair Module (NEB, E6050S) (79). C-tailing was performed on blunt-end DNA fragments using dCTP/ddCTP and Terminal Deoxynucleotidyl Transferase according to manufacturer guidance (Promega, M1871). Custom Tn RT fwd and Tn RT rev primers (Table S1) were used to PCR amplify C-tailed fragments to facilitate subsequent attachment of IDT for Illumina DNA/RNA UD Indexes (Illumina, Ref: 20026121, Lot: 20647691). Libraries were sequenced using NextSeq1000 (2×150bp) (Illumina, 20046813). Two biological replicates were sequenced per condition.

TnSeq data was pre-processed to remove transposon sequences using cutadapt (v3.4) (81) with default parameters from the R1 FASTQ file. After trimming, the sequencing data were aligned to the *Zymomonas mobilis* ZM4 genome (GCF_003054575.1-RS_2023_03_19) using bowtie (v1.3.1) (82) and default parameters. Transposon insertion sites were identified with TSAS (v2.0) (83) using the one sample analysis mode with minimum hits set to 5, clipping set to 5, capping set to 0, and weights set to 0. Essential genes in the absence of chemical treatment were classified by *p* < 0.05 and normalized unique hits per bp < 0.025 in the untreated and solvent control conditions (Table S7). Changes in fitness (i.e., chemical-gene scores) were identified using pairwise comparison in edgeR (v4.2.0) (84) to compare the number of insertions per gene in the experiment to the control samples.

### CRISPRi library sequencing and data analysis

Two biological replicate CRISPRi samples were sequenced per condition with the exception of the cinnamic acid conditions, which had one replicate per concentration. DNA was extracted from cell pellets using the Thermo Scientific GeneJET Genomic DNA Purification Kit (K0721), eluting in a final volume of 50 µL with typical yields ∼10-50 ng/µL. The sgRNA-encoding region was amplified by low-cycle Q5 PCR (19 cycles) as described previously (43), using primers oJMP697/oJMP698 or barcoded primers oJMP1678-1773/oJMP698 (85). PCR products were spin purified, and samples were sequenced by the UW-Madison Biotechnology Center Next Generation Sequencing Core facility using a NovaSeq 6000 (150 bp paired-end reads), at least 20 million reads per sample, and 20-30% PhiX when necessary for sequence diversity.

Pooled sample sequencing data were demultiplexed by primer barcode using cutadapt (v4.2) (81). As performed previously (43), sgRNA-encoding spacer sequences were counted using the seal.sh script (v38.90) from the BBTools package (https://sourceforge.net/projects/bbmap/), and counts were compared using edgeR (v.4.0.16) to calculate chemical-gene scores (log2 fold changes between treated and untreated conditions, LFC) and corresponding false discovery rates (84). Gene-level scores were calculated as the median LFC of perfect-match spacers targeting each gene, and gene-level significance was determined using the Stouffer’s P-value (poolr R package v1.1-1) based on the FDRs of the spacers (88).

Essential genes required for viability in the absence of chemical treatment were classified by gene-level significance ≤ 0.05 and median LFC ≤ −5.1 in the IPTG-induced final timepoint (T2_induced_) compared to the uninduced initial timepoint (T0_uninduced_) (Table S7). This median LFC cutoff was chosen to approximate a similar fitness cost to that used to define *Z. mobilis* essential genes previously (43). Since median LFCs in this study were generally stronger than those reported previously, likely due to longer outgrowth following induction, a linear regression (y = 1.567x - 0.373) was calculated representing the relationship between gene LFCs in each study. This regression was used to extrapolate the essential gene cutoff from Enright and Banta et al. (LFC ≤ −3) to the cutoff used in this work (LFC ≤ −5.1).

### Comparative TnSeq and CRISPRi analyses and visualization

Quantile normalization of TnSeq and CRISPRi chemical-gene scores was performed within each condition for all non-essential genes using the preprocessCore R package (version 1.64.0) (86). The pROC R package (version 1.18.5) was used to create receiver operating characteristic (ROC) curves and calculate area under the curve (AUC) (87). Heatmaps were generated using the ComplexHeatmap R package (version 2.18.0) (89). Hierarchical clustering was performed using Ward’s method. For essential *suf* genes, mismatched sgRNAs causing a partial fitness defect in the absence of chemical treatment, characterized by −4 ≤ LFC(T2_induced_/T0_uninduced_) ≤ −1 and FDR ≤ 0.05, were considered. The median chemical-gene score for these partial knockdown sgRNAs was calculated for visualization in heatmaps.

### Chrome Azurol S (CAS) Liquid Assay for Iron-Binding Compounds

The CAS assay protocol was modified from Wilhelm 2017 (version 2) (90). Chemicals dissolved in methanol were mixed 1:1 with CAS solution in a clear 96-well microplate and incubated in the dark for 1 hour at room temperature. Absorbance at 630 nm was measured using a Tecan Infinite M Nano plate reader. The relative chelation score was calculated as the OD_630_ ratio of methanol to chemical.

### β-Galactosidase assay

The *fhuE*::*lacZ* promotor fusion strain (PK17304) was grown at 30°C in ZRDM anaerobically via sparging (95% N_2_ and 5% CO_2_). At O.D._600_ 0.25, 2,2’-dipyridyl (DIP) or ferulic acid were added to a final concentration of 15 µM and 10 mM respectively. DMSO was included as a solvent control. Cell samples were collected for enzymatic assay prior toand 45 and 90 minutes after chemical treatment by adding chloramphenicol to a final concentration of 120 µg/mL to halt cell growth and protein synthesis before being stored on ice. β-galactosidase activity was then assayed to measure *fhuE* promoter activity (91).

### Proteomics sample growth and preparation

Strains were grown anaerobically in ZRDM via sparging to approximately O.D._600_ ∼0.3 before being treated with 1:100 v/v DMSO or 1M ferulic acid in DMSO (final concentration in culture 10mM). Each treatment had three replicates for each strain. Proteomics samples were harvested prior to and two hours after treatment by collecting 1mL culture, pelleting cells and freezing the cell pellet with dry ice before storing at −80°C.

Proteins were precipitated from *Z. mobilis* cells with 2:2:1 methanol:acetonitrile:H_2_O, then centrifuged at 12,000 x *g* for 10 minutes. Protein pellets were then dissolved in a lysis buffer containing reducing and alkylating agents (8 M urea, 100 mM Tris-HCl pH 8.0, 10 mM TCEP, 40 mM CAA). Protein concentration was quantified using BCA (Thermo) and brought to 1.5 mg/ml. Sequencing grade LysC (Wako) and trypsin (Promega) were added to the samples simultaneously, both at 50:1 protein:protease and samples were incubated overnight at room temperature. Samples were acidified with formic acid (FA) to pH 2 then desalted using Strata-X 33 µm Polymeric Reversed Phase cartridges. Peptides were then dried by speedvac and reconstituted in 0.2% FA, quantified by Nanodrop (A205), and brought to 1 mg/ml.

### Liquid Chromatography Mass Spectrometry-Based Proteomics and Data Analysis

For each sample, 500 ng of peptide was injected onto a 75 µm ID fused silica column packed (Shishkova 2018, PMID: 30179449) with 1.7 µm, 130 Å pore size, Bridged Ethylene Hybrid (BEH)-C18 particles (Waters) using a Vanquish Neo (Thermo Scientific). The column was maintained at 50°C inside an in-house made heater. Peptides were analyzed across a 74-minute separation where the mobile phase was ramped from 100% mobile phase A (0.2% FA) to 46% mobile phase B (80% acetonitrile / 0.2% FA / 19.8% H_2_O) before washing with 100% mobile phase B and then equilibrating with 100% mobile phase A.

Mass spectra were acquired using an Orbitrap Ascend tribrid mass spectrometer (Thermo Scientific) with data dependent acquisition (DDA). MS1 data was collected with the Orbitrap at a resolving power of 240,000 at 200 *m/z* from 300-1350 *m/z*, and an AGC target of 250%. With a cycle time of 1 second, MS2 data was collected on peptides of charge state 2-5 in the linear ion trap (scan rate Turbo) using advanced precursor determination (92) over a scan range of 150-1350 *m/z*, and 250% AGC with 12 ms max inject time. Monoisotopic precursor selection was used in peptide mode and dynamic exclusion was applied to ions for 20 seconds with a mass tolerance of +/−5 ppm. DDA proteomics data was analyzed with MSFragger (version: 22.0) using default settings and the *Z. mobilis* ZM4 reference genome (downloaded from Uniprot on 9/30/2024).

### Protein localization prediction

SignalP 6.0 was used to test the *Z. mobilis* proteome for proteins containing a Sec or Tat signal peptide indicative of cell envelope or extracellular localization (47). Proteins predicted to have a signal peptide with a likelihood greater than 0.9 were considered an envelope protein. To identify proteins which lack a signal sequence but have a transmembrane domain, the proteome was screened using DeepTMHMM 1.0 (48). Proteins which had either a signal peptide or a transmembrane domain were annotated as envelope proteins. All other proteins were categorized as cytoplasmic.

### STRING functional enrichment analysis

The STRING database (v12.0) (50) was used to identify functional patterns in within our proteomics data. For each strain, protein names and their corresponding log 2-fold change scores before and two hours after ferulic acid exposure were submitted to the Proteins with Values/Ranks feature on the STRING website (string-db.org). Local Network Clusters were used to identify patterns between strains.

## DATA AVAILABILITY

Raw sequencing data for TnSeq and CRISPRi samples is available through the NCBI BioProjects PRJNA1270032 and PRJNA1276497, respectively. Raw proteomics data is available in the MassIVE database (MSV000097906). Code used for this work is available at https://gitpub.wei.wisc.edu/alenright/orthogonal-chemgen-approaches-in-zmo.

## ACKNOWLEDGEMENTS

This material is based upon work supported by the Great Lakes Bioenergy Research Center, U.S. Department of Energy, Office of Science, Biological and Environmental Research Program under Award Number DE-SC0018409. A.L.E.S. was supported by the National Institute of General Medical Sciences of the National Institutes of Health under Award Number T32GM135066 and by the National Science Foundation Graduate Research Fellowship Program under Grant No. 2137424. Any opinions, findings, and conclusions or recommendations expressed in this material are those of the author(s) and do not necessarily reflect the views of the National Science Foundation.

We thank Dan Xie and Yaoping Zhang for preparation of *Z. mobilis* growth media, Bailey Bell and Tim Bugni for materials and guidance for CAS assays, and the University of Wisconsin Biotechnology Center for sequencing assistance. We thank other members of the Great Lakes Bioenergy Research Center, especially the Kiley, Peters, Sato, Hittinger, Coon and Computational Biology groups for shared research space, equipment, materials, and helpful feedback.

